# Short-term management of kelp forests for marine heatwaves requires planning

**DOI:** 10.1101/2025.01.30.634607

**Authors:** Jess K. Hopf, Anita Giraldo-Ospina, Jennifer E. Caselle, Kristy J. Kroeker, Mark H. Carr, Louis W. Botsford, Alan Hastings, J. Wilson White

## Abstract

Heatwaves are now pervasive stressors to marine ecosystems, and it is urgent to consider mitigation tools that support ecosystem resilience and persistence in the immediate future. We modeled a system of kelp, herbivorous urchin, and predatory fish to compare how potential management actions (kelp seeding, urchin removal, and fishery closures) could reduce the likelihood of a heatwave shifting a kelp forest into a degraded urchin barren state. We found that those interventions were most effective when begun alongside or before the start of a heatwave. Closing the predatory fish fishery was more effective when done earlier and for longer, while urchin removal and kelp seeding were more effective when begun alongside and continued throughout the heatwave. Kelp seeding was notably less effective than other interventions. Our results suggest the need for improved heatwave forecasting and nimble management protocols to enact mitigation actions quickly if a heatwave is forecasted or occurs.

## Introduction

The acceleration of climate change has come with an urgent need to understand what local actions can mitigate impacts. In particular, the increased frequency, intensity and duration of marine heatwaves (hereafter ‘heatwave(s)’) are a threat to marine systems and the ecosystem services they provide (Frölicher et al. 2018, Smith et al. 2021, 2023). Heatwaves can be defined as discrete pulses of extreme ocean temperatures occurring against the background of chronic warming (Hobday et al. 2016, Amaya et al. 2023). While long-term management responses are required to address long-term temperature trends, short-term, temporary interventions (our focus here) are critical to support systems through discrete heatwave events. The key questions are what management actions will be most effective, and when to implement them (Ainsworth et al. 2020, Pinsky et al. 2021, Smith et al. 2021, 2023).

Heatwaves can stress certain species while benefiting others, resulting in complex ecosystem responses. In northeastern Pacific kelp forests, warm, low-nutrient conditions during heatwaves reduce growth and recruitment of the dominant species, giant kelp (*Macrocystis pyrifera*) (Zimmerman and Kremer 1986, Hollarsmith et al. 2020, Michaud et al. 2022). Conversely, sea urchins (key herbivores; *Mesocentrotus franciscanus* and *Strongylocentrotus purpuratus*) experience increased grazing rates (Spindel 2023). The decrease in standing kelp also reduces production of detached kelp fronds (‘drift kelp’), a preferred food source for urchins (Kriegisch et al. 2019, Randell 2022). The loss of drift kelp combined with increased grazing rates can trigger urchins to switch behavior, from cryptic feeding on drift and detritus in rocky crevices to exposed roaming and targeting standing kelp stipes (Kriegisch et al. 2019, Smith and Tinker 2022, Rennick et al. 2022; Figure 1). As a result, heatwaves can sometimes tip the ecosystem from a kelp-dominated forest to an urchin barren (McPherson et al. 2021). Recovery from the degraded barren state can be challenging even after returning to kelp-beneficial conditions (i.e., the system exhibits hysteresis; Filbee-Dexter and Scheibling 2014, Ling et al. 2015) largely because the barren-state urchins have high survival, despite low food availability and exposure to predators. In fact, some predators actively avoid starved, barren-state urchins (Eurich et al. 2014, Liebergesell 2022, Smith and Tinker 2022). Thus, kelp resilience to heatwaves is a fine balance of consumer-resource dynamics.

**Figure 1:**
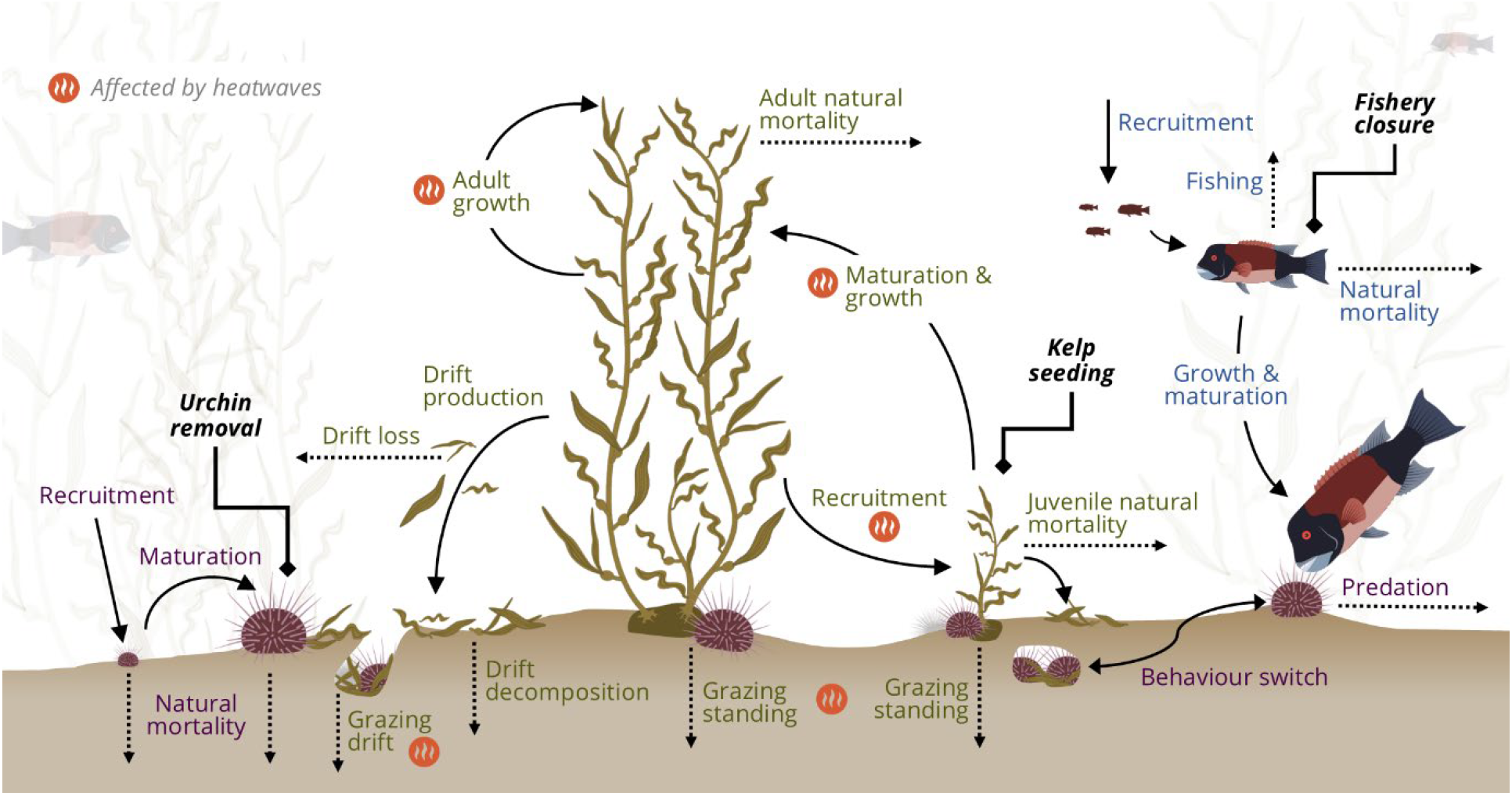
Three-species model overview. Main processes included in our kelp-urchin-predator model, including those affected by heatwaves (orange symbol). Purple and green text show processes relevant to the kelp-urchin sub-model, and blue text show processes relevant to the predator sub-model. Bold, black text indicates potential mitigation actions considered in this study. All artwork by Jess K. Hopf, except giant kelp icons, which are modified from artwork by Jane Thomas sourced from ian.umces.edu/media-library under the CC BY-SA 4.0 license.

Kelp forests are ecologically and economically important, and we need to understand how to mitigate heatwave effects on them (McPherson et al. 2021, Hamilton et al. 2022, Eger et al. 2023, CDFW 2023). Management actions can be broadly grouped by which link in the kelp-urchin-predator trophic chain they address (Figure 1). First, direct kelp restoration aims to increase kelp biomass via seeding spores or transplanting sporophytes (Eger et al. 2022). Second, herbivore removal can promote kelp recovery by removing the key herbivore(s) via harvest, culling, or translocation (Eger et al. 2022).

Third, because many urchin predators (e.g., fishes, lobsters, sea otters) have declined due to human harvest, increasing predator densities through fishery closures can suppress herbivory and support kelp forest persistence (Eisaguirre et al. 2020, Kumagai et al. 2024).

Here we focus on potential short-term management actions: temporary changes enacted in response to an impending, current, or recently-ended heatwave that may prevent a shift from a kelp forest to an urchin barren (i.e., ecosystem resistance). We help address the management question, ‘what action(s) could be taken in the face of a heatwave and when?’ This reflects a proactive or reactive response to sudden, acute changes in the system rather than permanent management actions that would better orient the system towards longer-term goals (Pinsky et al. 2021, Arroyo-Esquivel et al. 2023). To do this, we analyzed a three-species model that captures the interactions among giant kelp, purple urchins (*Strongylocentrotus purpuratus*), and a fishery-targeted urchin predator (Californian Sheephead; *Bodianus pulcher*, henceforth ‘Sheephead’) (Figure 1). We use Southern Californian (USA) kelp forests as our study system, but our results are applicable to kelp systems globally where urchin predators are targeted by fisheries. We consider the ability of three main management actions (kelp seeding, urchin removal, and a temporary predator fishery closure) to reduce the likelihood of a kelp forest shifting to an urchin barren because of a heatwave when the actions are implemented before, during, or after a 2-year heatwave. This analysis complements studies that consider these management actions only after forests have shifted to barrens states (e.g., Dunn et al. 2021, Arroyo-Esquivel et al. 2023).

## Methods

### Three-species population model

We developed a three-species, single patch (spatially-implicit) population model that captures the interactions between giant kelp, herbivorous purple urchins, and a fishery-targeted predatory fish (Californian Sheephead) (Figure 1). The model is comprised of two sub-models that run over time: 1) a kelp-urchin stage-based sub-model that tracks biomasses of key life-stages: juvenile and adult standing kelp, drift kelp, juvenile urchins, and exposed and hiding adult urchins, and 2) a predatory fish integral-projection sub-model (IPM), which tracks the abundance of sheephead of all lengths over time; we then translate length-abundance into biomass. To account for seasonal variations in ecological processes (recruitment, kelp growth, urchin grazing, and heatwave effects), the model uses seasonal (3-month) time steps. Details are provided in the Supplemental Information, and all code is publicly available at repository DOI:0.5281/zenodo.15606579.

Our model captured the characteristic kelp forest dynamic in which overgrazing by urchins shifted the system from a kelp forest to an urchin barren with local kelp extinction. Underpinning this was an urchin behavioral feeding switch: when drift kelp was abundant, urchins fed passively in crevices on drift, but left crevices to actively graze on standing kelp when drift kelp was scarce. To achieve this, we modelled the proportion of exposed urchins as a declining function of drift biomass density relative to urchin consumptive capacity (Rennick et al. 2022), building on Randell’s (2022) approach. We assumed a Type-II functional response for kelp grazing by urchins, and a Type-I response for urchin predation by Sheephead, as a function of the total biomass of Sheephead large enough (>20 cm; Hamilton and Caselle 2015) to consume adult urchins. Exposed urchins had a higher rate of predation mortality than cryptic urchins (Nichols et al. 2015). However, if there was no kelp for >3 months, then Sheephead stopped consuming urchins, reflecting the reduced nutritional value of starved urchins (Liebergesell 2022). We assume a spatial domain equivalent to a spatially isolated kelp forest, at which scale the Sheephead and urchin populations would be open with external recruitment, and the kelp population would be largely closed (1-10 ha; Kinlan and Gaines 2003), with recruitment a function of local adult biomass and density dependence through shading and competition (Nisbet and Bence 1989). This can be viewed as a worst-case scenario for kelp resilience without rescue effects from external kelp populations. Stochasticity was captured through year-to-year variation in recruitment for all species, with recruitment each year drawn randomly from data-parametrized normal distributions.

### Modelling heatwaves & management strategies

All modelled scenarios began in an undisturbed kelp forest state. Reflecting the known effects of heatwaves on kelp and urchins, we implemented a heatwave as a period of reduced kelp recruitment (Hollarsmith et al. 2020) and growth (Zimmerman and Kremer 1986), and increased urchin grazing rates (Spindel 2023). The modelled heatwave started in winter and lasted 2 years, following the recent 2014-2016 Californian heatwave (Michaud et al. 2022).

We considered three possible management actions: a temporary Sheephead fishery closure, urchin removal (a set biomass of adult urchins removed each season), and kelp seeding (a set biomass of juvenile kelp added each season). We focused on kelp seeding, rather than transplantation of adult kelp, as seeding is more viable on a large scale (Eger et al. 2022). We modelled scenarios where actions began up to two years before the heatwave (i.e., pre-emptive actions; while seeding into an extant kelp forest is unlikely, we considered that possible action for the sake of completeness), during the heatwave, or up to three years after the heatwave ended, and actions lasted one to five years. We also explored a range of magnitudes for urchin removal and kelp seeding. Finally, we considered the implications of simultaneously implementing combinations of two management actions.

We simulated 10^5^ replicates of each scenario, running the model for 20 years prior to the heatwave and then assessing the proportion of simulations in which a kelp forest persisted (defined as non-zero kelp biomass for at least a year) 8 years after the heatwave ceased (enough time to complete all potential management actions). Monitoring sooner only reduced the realized impact of a temporary fishery closure (Figure S2.3; Supplemental Information). We present this response variable as *the percentage increase in probability of kelp forest persistence* relative to the effect of the heatwave with no management intervention, and to a baseline scenario with no heatwave (Figure 2), such that:

**Figure 2:**
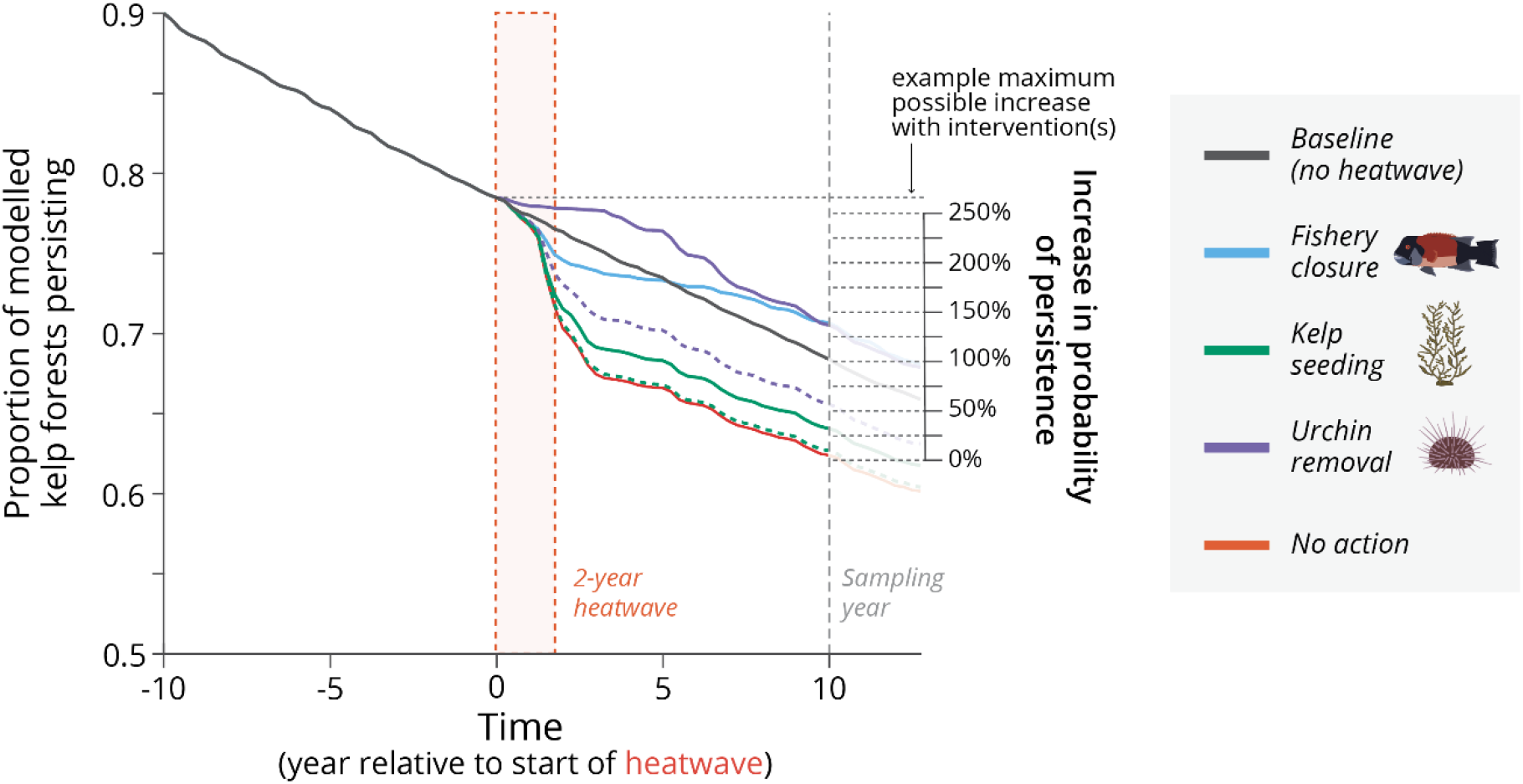
Example scenario runs. The proportion of model simulations (n = 10,000) with kelp forests persisting over time, and our response variable, *the percentage increase in probability of kelp forest persistence*, calculated at the sampling year (eight years post-heatwave), for example scenario runs. In this example, the mitigation actions began the same year as the heatwave started, and lasted three years. For urchin removal and kelp seeding, 10% (dashed lines) and 100% (solid lines) of the pre-heatwave biomass was removed/added yearly. Mitigation actions can only increase the probability of persistence to a maximum point, equal to that at the start of the intervention.

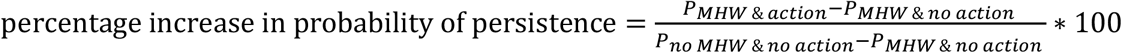

where *P*_*i*_ is the proportion of simulations with persistent kelp forests under scenario *i*. Note that even without a heatwave, the proportion of kelp forests persisting declined over time (Figure 2) because in a stochastic system (and without kelp or drift immigration) there is a small, constant probability that an individual forest will switch to a barren in any given time step due to a series of poor kelp, and/or substantial urchin recruitment events.

## Results

The three standalone management actions considered were most effective at increasing the probability of a kelp forest persisting through the heatwave if initiated prior to, or during the first year of the heatwave (Figures 3-4). Notably, kelp seeding was markedly less effective overall than urchin removal or fishery closures. Interventions were still moderately effective (>25% increase in probability of persistence) after the heatwave has ended, but only if undertaken intensively for 3 or more years (Figure 3). This was generally robust to changes in key parameter values (based on sensitivity analyses; see Supplemental Information).

**Figure 3:**
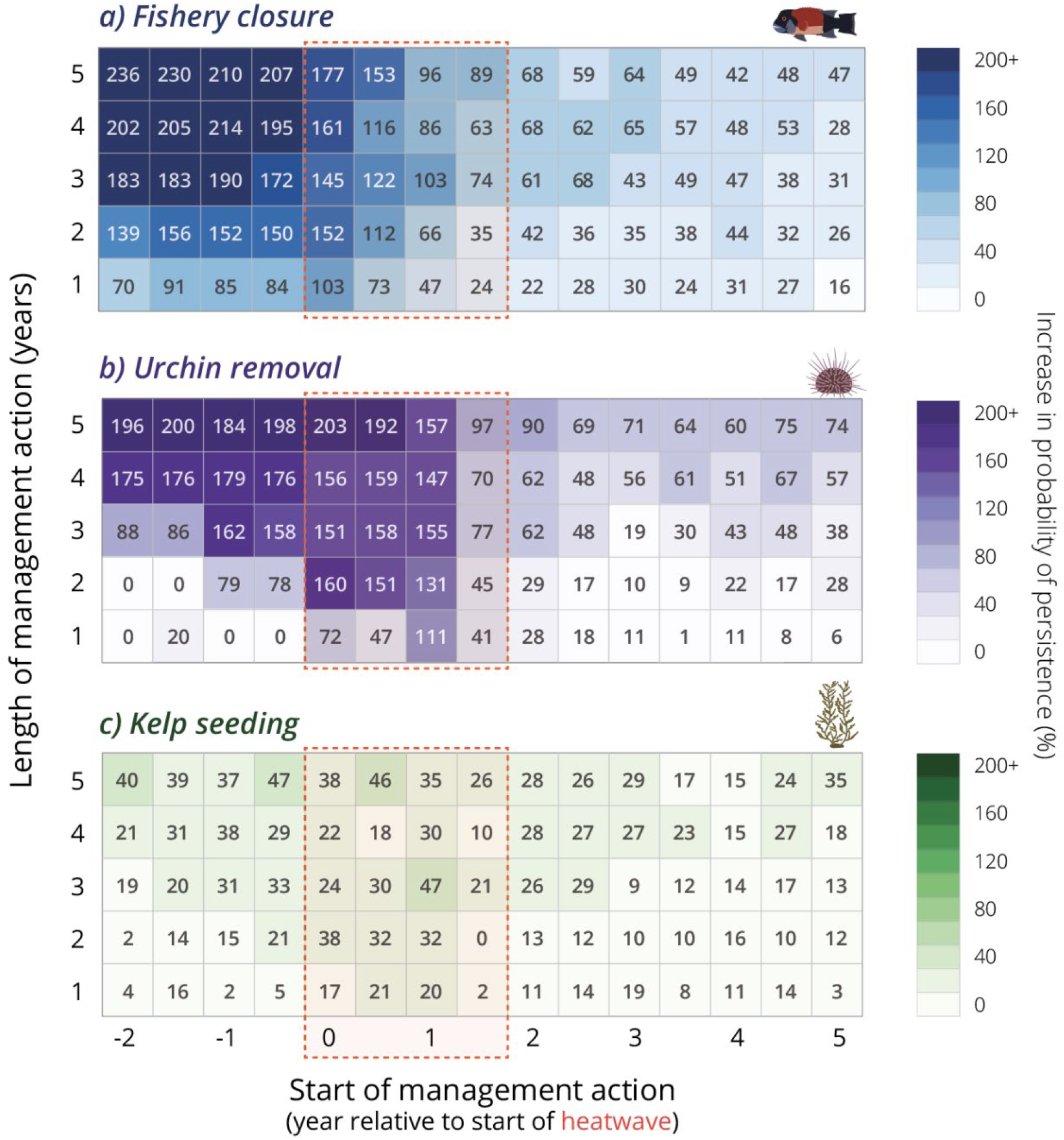
Efficacy of individual heatwave mitigation actions. Percentage increase in the probability of kelp forest persistence through a 2-year marine heatwave with individual management actions, over length and timing of action. These modelled scenarios present the most intensive management scenario in which 100% of the pre-heatwave urchin and kelp biomass is removed or seeded each year, respectively. The orange dashed box indicates the action scenarios that began during the heatwave. Note that all actions are enacted temporarily.

Fishery closures were more effective the earlier and longer they were implemented: for example, closures that began 6+ months before the heatwave and lasted a minimum of 3 years were extremely effective (>200% increase in the relative probability of kelp persistence) (Figure 3a). In contrast, urchin removal and kelp seeding needed to overlap completely with the heatwave to be most effective (Figure 3b-c, Figure 4). Both kelp and urchin actions were markedly less effective if completed before the heatwave began (bottom left of Figure 3b-c).

**Figure 4:**
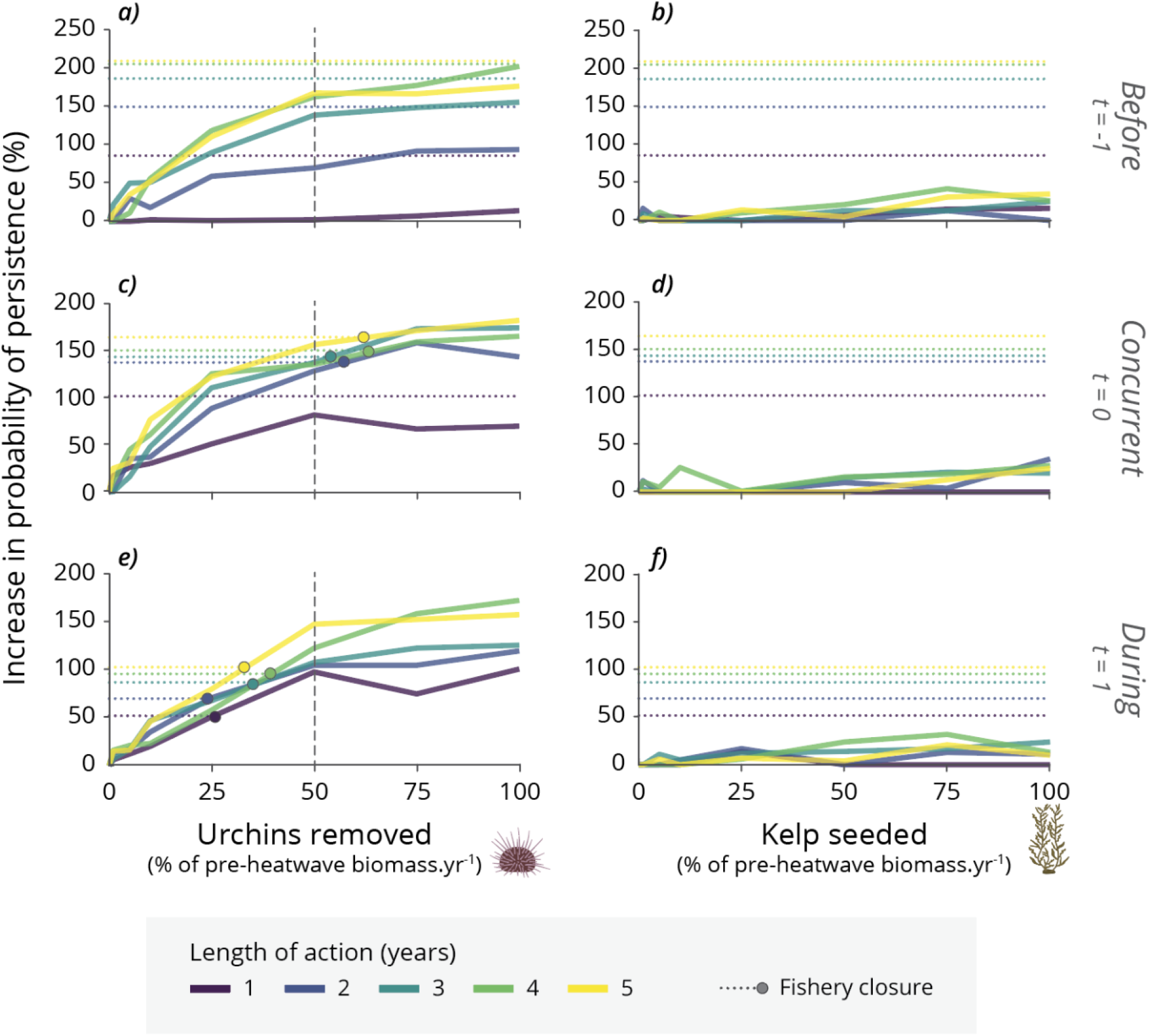
Changes in efficacy with effort. Percentage increase in the probability of kelp forest persistence through a 2-year marine heatwave with varying degrees of urchin removal (a, c, e,) and keep seeding efforts (b, d, f), and lengths of actions (colors), at different times (*t*, in years) relative to the start of the heatwave (rows). Dotted lines with points indicate efficacy for fishery closures starting at the same time; lines without points indicate that temporary fishery closures were more effective than 100% kelp seeding or urchin removal efforts. Degrees of effort are measured as the percentage of pre-heatwave urchin and kelp biomass removed or seeded each year, respectively.

While longer and earlier fishery closures were more beneficial, the same was not true for urchin removal and kelp seeding (Figures 3-4). So long as urchin removal and kelp seeding spanned the heatwave, their effectiveness varied little with timing or duration (Figure 3-4). For example, a 2-year action begun at the onset of the 2-year heatwave and a 4-year action begun 2 years prior were comparably effective (Figure 3). Figure 3 presents the most intensive management scenario in which 100% of the pre-heatwave urchin and kelp biomass is removed or seeded each year, respectively.

However, that intense effort was not necessary for urchin removal: there was little additional benefit to removing more than 50% of the pre-heatwave urchin biomass each season regardless of timing or length of the management action (vertical dashed lines; Figure 4a,c,e). The effectiveness of kelp seeding, although remaining low, generally increased with increasing effort (Figure 4b,d,f).

Reflecting the different optimal timings for the two most effective management interventions, fishery closures that began a year before the heatwave were generally more effective than urchin removal implemented at the same time (Figure 4a). As the heatwave progressed, the relative effectiveness of fishery closures and urchin removal shifted so that, one year into the heatwave, fishery closure became less effective then removing at least 25-40% of the pre-heatwave urchin biomass were more effective than fishery closures (dotted lines and dots in Figure 4c,e).

Combining management actions of varying degrees had additive effects, at best, but typically reached a threshold (Figure 1) beyond which additional management efforts provided little benefit (Figure 5). We demonstrate this by focusing on the scenario where 3-yr long actions are implemented at the start of the heatwave. The highest gains for combing actions occurred when combining moderate degrees of urchin removal (>50%) with fishery closures (Figure 5). Given the low efficacy of kelp seeding relative to urchin removal or fishery closures, little benefit was gained from combing kelp actions with other interventions.

**Figure 5:**
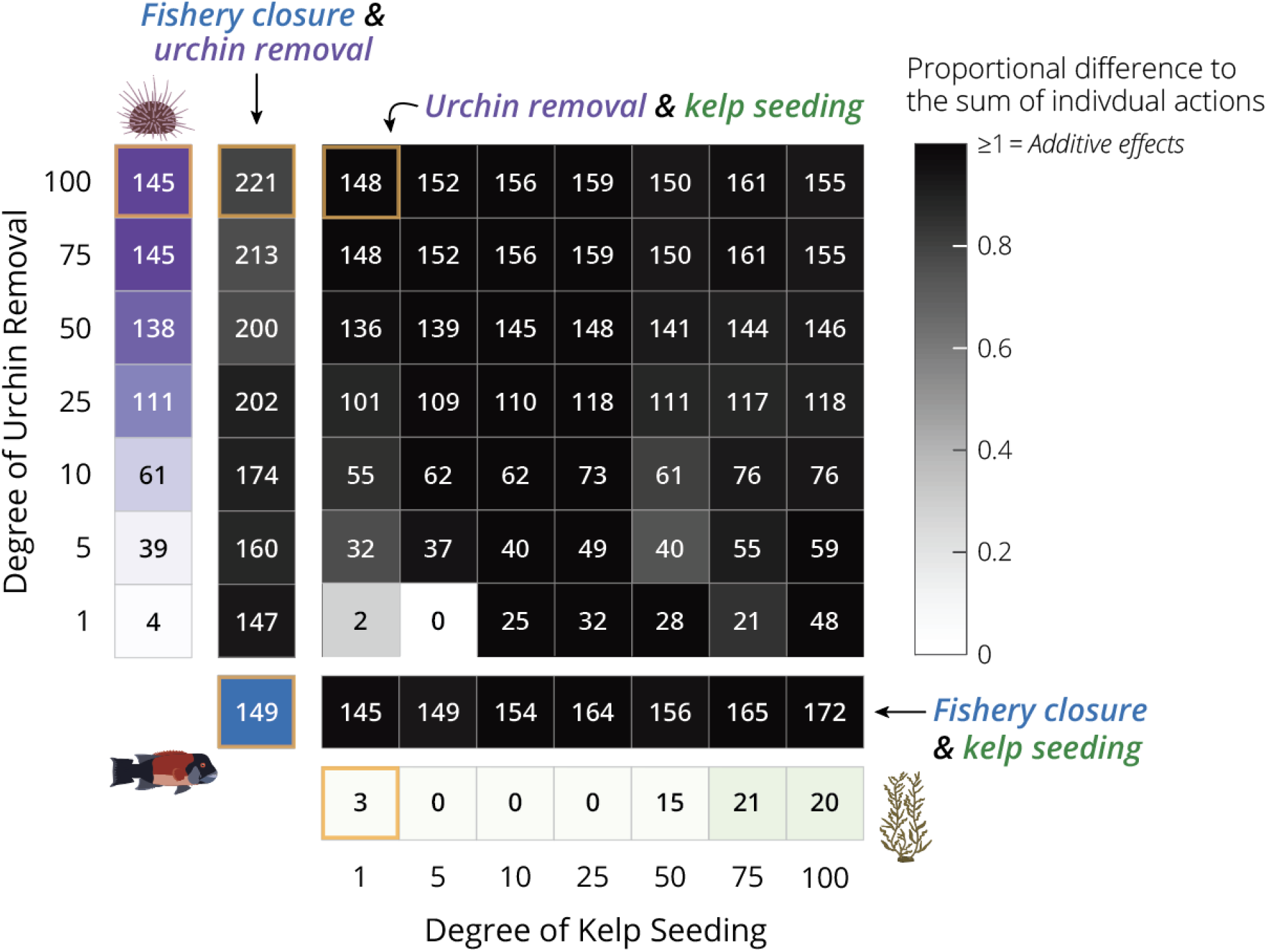
Efficacy of combining two heatwave mitigation actions. Modelled percentage increases (±10 units) in the probability of a kelp forest persisting through a 2-year marine heatwave (numbers in cells), and the proportional difference between the efficacy outcome of the combined actions compared to the expected summed outcome of the actions based on their individual efficacy (grey color scale), over a range of urchin removal and kelp seeding efforts. In this demonstrative scenario, actions occur simultaneously at the start of the heatwave and last for the same period of time (3 years). Blue, purple, and green boxes show the efficacy of individual actions for comparison for Sheephead, urchins and kelp respectively (see also Figure 3). As an example (boxes outlined in orange), in the top left corner of the figure, adding 1% kelp seeding and 100% urchin removal has an *additive* effect (3% effect of kelp alone + 145% effect of urchin alone ≈ 148%, the combined effect of both) but 100% urchin removal plus a fishery closure is *less than additive* (145% effect of urchin alone + 149% effect of fish alone > 221%, the combined effect). The degree of urchin removal and kelp seeding effort is the percentage of pre-heatwave urchin and kelp biomass removed or seeded each year, respectively. Note that all actions are enacted temporarily.

## Discussion

Short-term, temporary management actions may be essential to supporting kelp forests through heatwaves. We found that timing is critical: interventions begun before a heatwave (for temporary fishery closures) or during the first year of a heatwave (for urchin removal and kelp seeding) were most effective in reducing the probability that a kelp forest would shift to an urchin barren (measured 8 years after the heatwave ended; but see also Supplementary Information). As heatwave forecasts are currently available only months beforehand at best (Holbrook et al. 2020, Jacox et al. 2022), our results suggest that action plans (e.g., protocols, permits, and funding) need to be in place preemptively, so that mitigation actions can be enacted quickly. This also highlights the need to continue to develop longer, more robust forecast warnings so that effective actions can be implemented in time (Holbrook et al. 2020, Jacox et al. 2022). As forecasted heatwaves are not a certainty, cost-benefit analyses are needed to characterize potential trade-offs between the costs of undertaking an action that was not needed (if a potential heatwave does not occur) and failing to undertake an action that was needed. Such analyses need to consider the logistical and monetary costs as well as the local management goals and social acceptance of different actions, which should be assessed on a case-by-case basis.

Like others (Dunn et al. 2021, Arroyo-Esquivel et al. 2023), our findings reinforce the need to consider underlying ecological processes in management plans. To buffer against kelp forest collapse, fishery closures require years (depending on the life history of the species; Kaplan et al. 2019) to rebuild predator biomass to levels sufficient to suppress urchin densities and prevent overgrazing during a heatwave. Therefore, we found longer and earlier predator fishery closures more effective, such as could be achieved with a permanent no-take marine reserve (White et al. 2025). As preemptive management actions are unlikely, or potentially too costly/risky, to occur before a heatwave starts, and highly unlikely to occur earlier than 1 year prior to a heatwave (due to forecast limitations), the full effectiveness of a temporary fishery closure may be limited, especially compared to large-scale, highly effective urchin removal efforts. Conversely, temporary fishery closures that start at the beginning of the heatwave can be as effective as low-to-moderate urchin removal.

Managers would need to consider the costs, trade-offs, and feasibility at different spatial scales, of those alternatives when assessing options. Opening urchin fisheries could also act as an urchin removal action, although cost-benefit analyses would need to account for poor gonad quality in urchin barrens. Predator fishery adjustments (e.g., quotas, closures, and restocking) have already been implemented in response to heatwaves, although often retrospectively and with maintaining stock biomass rather than ecosystem persistence in mind (Smith et al. 2021). Nimble ecosystem-based fishery management could focus on regulations that maintain the desired level of urchin consumption by predators; the full closures we modeled could be viewed as a precautionary ecosystem-based intervention in cases where the amount of urchin predation needed for ecosystem persistence is uncertain.

In contrast to fishery closures, removing urchins or seeding juvenile kelp are relatively quick-acting but short-lasting, logistically costly, and challenging at scale (Eger et al. 2020b, 2022). Kelp seeding, in particular, had limited success in our model, due to density-dependent feedbacks (here, we included shading of juveniles by the established kelp canopy, and competition between recruits; Nisbet and Bence 1989, Reed 1990) and urchin grazing. Although our model included the effects of reduced kelp recruitment and growth during warmer temperatures, we assumed that initial seeded kelp establishment was always successful, even during heatwaves, while the subsequent growth of restored sporophytes was heatwave affected. Less successful seeding would be equivalent to lower efforts in Figure 4. Our results also show that urchin and kelp interventions require ongoing intervention throughout the heatwave to maximize effectiveness. For example, external larval supply for urchins can drive fast rebounds following removal, necessitating continuous removal attempts through the heatwave. This effect would be exacerbated by immigrating adults, which we have assumed is negligible at the scale of our model. Likewise, starting these actions earlier may waste valuable resources, as we found that urchin removal and kelp seeding actions that occurred prior to and continued through heatwave were as effective as those that start the same year as the heatwave.

Interestingly, we found that combining two interventions did not always increase effectiveness as much as could be assumed based on their individual outcomes, especially with fishery closures and large urchin removal efforts. The implications of this will depend on the per-kilogram cost of action, but it suggests that there may be benefits to combining actions if increasing efforts of an individual action yields diminishing returns, compared to undertaking two actions. There may also be benefits to offsetting actions in time, such as an early fishery closure followed by urchin removal. However, testing all such possible combinations was outside the scope of this study. We recommend that case-specific modelling and cost-benefits analysis be undertaken.

A key assumption of our model was a closed kelp population without larval rescue effects or external supplies of drift kelp. We also assumed a constant influx of urchin recruitment, independent of the local urchin densities. These assumptions consider the worst-case scenario for kelp resilience, and drive the observed low success of post-heatwave management actions. However, we expect that our general findings will still hold true for areas that do experience notable external kelp recruitment, as such recruitment events are patchy in space and time (Cavanaugh et al. 2013) and should not be relied upon, although they may benefit kelp-forest restoration efforts (Eger et al. 2020a). Lastly, we focused on Sheephead, but our results would be applicable to other fishery-targeted predators like the Spiny Lobster (*Panulirus interruptus*) or Southern Rock Lobster (*Jasus edwardsii*), which are also mobile benthic invertivores that suppress kelp herbivory when protected from harvest (e.g., Peleg et al. 2023).

Many climate-mitigation options are being explored for a range of marine systems (e.g., Ainsworth et al. 2020), with increasing focus on optimizing these actions for success (e.g., through site-selection, or synergistic actions; Eger et al. 2020a, Brown et al. 2024). Our study demonstrates that the timing of a mitigation action, relative to a heatwave, critically affects its success in reducing the likelihood of a kelp-forest to urchin-barren shift. Most heatwave responses to date have been reactive (Smith et al. 2021) and our results suggest that managers should begin to also consider analyzing options and establishing protocols for preemptive heatwave mitigation actions.

## Acknowledgements

We thank Pete Raimondi, Mark Novak, Andrés Pinos-Sánchez, and Iris Flores for helpful thoughts during model development and analysis. J. K. Hopf thanks Cashie the wolfhound-cross for her supportive snoring during the writing of this manuscript. This work was funded by the David and Lucile Packard Foundation (award 2022-74730), the California Ocean Protection Council (agreement C0874012), and Oregon Department of Fish and Wildlife (contract IGA 251-23). This is publication XX of the Partnership for Interdisciplinary Study of Coastal Oceans, funded primarily by the David and Lucile Packard Foundation.

## Notes

**Conflict of interest disclosure:** The authors have no conflicts of interest to declare.

### Competing Interest Statement

The authors have declared no competing interest.

### Summary of Updates

Updated methods to capture more realistic density-dependent survival in kelp and new results.

