## Supplementary materials for "Short-term management of kelp forests for marine heatwaves requires planning"

#### *Short-term management of kelp forests for marine heatwaves requires pre-emptive action*

1. Supplemental Methods 3
1.1. Model overview 3
1.2. Sheephead sub-model 3
1.3. Urchin-kelp sub-model: purple urchins 6
1.4. Urchin-kelp sub-model: giant kelp 9
2. Sensitivity analyses 17
3. References 23

### 1. Supplemental Methods

*1.1. Model overview*

We developed a three-species, single patch (spatially-implicit) population model that captures the interactions between giant kelp (*Macrocystis pyrifera*; hereafter 'kelp'), herbivorous purple urchins (*Strongylocentrotus purpuratus*), and the fishery-targeted, urchin-predating Californian Sheephead (*Bodianus pulcher*). The model comprises of two sub-models that run over time: 1) a predatory fish integral-projection sub-model (IPM; following Hopf and White 2023), which tracks the abundance of sheephead of all lengths over time; we then translate length-abundance into biomass using an allometric relationship; and 2) an urchin-kelp stage-based sub-model that tracks biomasses of key life-stages: juvenile and adult standing kelp, drift kelp (detached kelp fronds), juvenile urchins, and exposed and hiding adult urchins.

The model runs over seasonal (3-monthly) time steps, following the Northern Hemisphere seasons: winter (December – February), spring (March – May), summer (June – August), and autumn (September – November).

Where possible, the model is parameterized using data from the Channel Islands, southern California, otherwise data from comparable regions are used (see Tables S1.1 & S1.3 for specifics). Data sources are either published literature (see Tables S1.1 & S1.3) or published datasets (PISCO; access through DataONE). Models were run in MATLAB (The MathWorks Inc. 2022). All code can be found at DOI: 10.5281/zenodo.15606579.

*1.2. Sheephead sub-model*

To model Sheephead population dynamics over time, we used an integral-projection model (IPM; Ellner and Rees 2006, Rees et al. 2014, White et al. 2016), which models dynamics in a size-structured, discrete-time manner. This model structure captures well changes in population size structures over time, which translates into changes in predation of urchins, as larger Sheephead consume more urchin biomass. Our model structure closely follows Hopf and White (2023). All model parameter descriptions and values for the Sheephead sub-model are in Table S1.1.

The Sheephead IPM tracks the state of a population in terms of its size distribution, $n(x,t)$, which is the density function of individuals of size $x$ in season $t$. The population density at time $t + 1$ is the current population density multiplied by the probability density of moving from size $x$ to size $y$, $K(y,x)$, plus new recruits, integrated over all biologically reasonable sizes $\Omega$:

$$n(y,t+1) = \int_{\Omega} K(y,x)\, n(x,t)dx + R_S\, \rho(y)\, \theta_S.$$

The kernel $K(y,x)$ is a product of the growth kernel for survivors, $G(y,x)$ (a von Bertalanffy growth function), and survivorship, $P(y,x)$, which is size-dependent:

$$P(y,x) = e^{-[M-\phi(x)F]}G(y,x)\ ,$$

where $M$ is instantaneous natural mortality rate per season, $F$ is the fishing mortality rate per season, and $\phi(x)$ is the selectivity function describing the vulnerability of size $x$ fish to harvest. The selectivity function $\phi(x)$ is zero below the legal size-limit and one above it.

Assuming an open population structure for Sheephead, recruitment is added as the product of the probability density function for initial recruit size $\rho(y)$ and density of new recruits arriving $\overline{R_S}$, which varies year-to-year. $\theta_S$ is the recruitment timing function for Sheephead, which captures autumn peaks in Sheephead recruitment (Cowen 1990, 1991, Alonzo et al. 2004). Note that each year in the model, total *yearly* recruit abundance is drawn from a normal distribution ( $norm(\mu_F, \sigma_F)$ ) and $\theta_S$ distributes this recruitment proportionally over the seasons and sums to one each year. Increased Sheephead recruitment was noted during the 2014-2016 Northeast Pacific marine heatwave (Kumagai et al. 2024), however we did not include this as a heatwave effect as it unclear whether the observed event was due to increased temperatures or altered oceanographic conditions. As the effects of temporary increased Sheephead recruitment would be consistent across all scenarios, we do not expect this effect would alter our results.

The total biomass of Sheephead predating urchins in a given season $t$ ($N_t$) is therefore the product of the abundance density function, $n(y,t)$, and the weight-at-length function, $W(y)$, integrated for all sizes greater than then average size of Sheephead that are able to effectively consume urchins, $a_{pred}$:

$$N_t = \int_{x_{pred}}^{\infty} n(y,t)\, W(y)\, dy.$$

$W(y) = g\ y^h$, where $g$ and $h$ are functional shape parameters.

To remove the effects of transient dynamics in predator biomass densities, the first 40 years (long enough for transient dynamics to settle) of the Sheephead model were discarded. We assume that the fishery has been in place long enough that the fished population has reached a time-averaged equilibrium.

**Table S1.1** Parameter descriptions, values, and references (where applicable) for the Sheephead sub-model.

| Symbol | Description | Value | Reference/Source |
|---|---|---|---|
| $M$ | Adult instantaneous natural mortality rate (per season) | 0.2/4 | Alonzo et al. (2004) |
| $F$ | Adult instantaneous mortality rate due to fishing (per season) | 0.2/4 | Alonzo et al. (2004) |
| $x_F$ | Legal size limit | 30 cm | Alonzo et al. (2004) |
| $\boldsymbol{x_{pred}}$ | Minimum fish size able to effectively consume urchins | 20 cm | Hamilton and Caselle (2015)<br>Selden et al. (2017a) |
| $\boldsymbol{R_S}$ | Yearly density of new Sheephead recruits (including variability)<br>$\mu_F$ = mean<br>$\sigma_F$ = standard deviation | <br>355<br>0.7188 | Mean value set to achieve average unfished of Sheephead densities in Channel Islands kelp forests, based on PISCO data*.<br>Variance based on PISCO data* |
| $\boldsymbol{\theta_S}$ | Sheephead recruitment timing function | [0 0 0.05 0.95]$^{+}$ | Cowen (1990, 1991)<br>Alonzo et al. (2004) |
| $g$ | Weight-at-length shape parameter | $2.7\text{x}10^{-5}$ | Alonzo et al. (2004) |
| $h$ | Weight-at-length shape parameter | 2.86 | Alonzo et al. (2004) |

* See PISCO_kelp-urchin-Sheephead_data.Rmd file for details.

$^{+}$ Seasonal variations, in the order winter, spring, summer, autumn

*1.3. Urchin-kelp sub-model: purple urchins*

See Table S1.2 for state variables, and Table S1.3 for model parameters descriptions and values for the urchin-kelp sub-model.

As we are including a kelp-drift feeding switch in urchin behavior, we chose a stage-based matrix model for the kelp-urchin sub-model, which easily allows attribution of different demographic rates to different life-history or behavioral stages. Following a stage-based matrix approach (Caswell 2001), the biomass density of urchins in the next time-step ($\mathbf{U_{t+1}}$) is a product of the biomass of urchins in the current time step and the transition matrix ($\mathbf{M_u}$), plus recruits ($\mathbf{R_{U,t}}$), such that

$$\mathbf{U}_{t+1} = \mathbf{M_{U,t}} * \begin{bmatrix} u_{J,t} \\ u_{C,t} \\ u_{E,t} \end{bmatrix} + \begin{bmatrix} \mathbf{R_{U,t}} \\ 0 \\ 0 \end{bmatrix},$$

where *J*, *C*, and *E* are the urchin life-stages: juveniles, cryptic adults, and exposed adults, respectively.

For urchins, we assume open-patch dynamics representing the case that the urchin patch is a portion of a larger population. This is a fair assumption at the scale of our model (1 ha patch) as urchins have long larval durations and distances (Kinlan and Gaines 2003, Arafeh-Dalmau et al. 2022). Urchin recruitment varies seasonally, with ~90% of recruitment occurring between March to July (spring into summer; Okamoto et al. 2020a), and shows interannual variation (Ebert et al. 1994). Therefore, the biomass of incoming urchin recruits ($\mathbf{R_{U,t}}$) is

$$\mathbf{R_{U,t}} = norm(\overline{R_U}, \sigma_{RU})\, \theta_{U,t}\, (s_J)^{\tau},$$

where $\overline{R_U}$ is yearly mean biomass and $\sigma_{RU}$ is the yearly variance of incoming urchin recruits, and $\theta_{U,t}$ is the timing function for urchin recruitment (capturing seasonal variation). Note that each year in the model, total *yearly* recruit biomass is drawn from a normal distribution, and $\theta_{U,t}$ distributes this recruitment proportionally over the seasons and sums to one each year. This is because empirical estimates of recruitment are yearly. Since planktonic larval durations (PLDs) for urchins are typically less than 3 months (40-90 days; Arafeh-Dalmau et al. 2022), recruits will operationally spend a portion of time as juveniles during a time step. We therefore included, in the recruitment function, a juvenile survival term ($s_J$) that is discounted by the proportion of time spent as a recruited juvenile ($\tau$): $\tau = 1 - \frac{PLD}{91}$. Note, that when $\tau = 1$, $(s_J)^{\tau} = s_J$.

The stage transition matrix ($\mathbf{M_u}$) for urchins is

$$\mathbf{M_{U,t}} = \begin{array}{r} \\ \textit{juvenile} \\ \textit{cryptic} \\ \textit{exposed} \end{array} \begin{array}{c} \begin{array}{ccc} \textit{juvenile} & \textit{cryptic} & \textit{exposed} \end{array} \\ \begin{bmatrix} (1-g_U)\, s_J & 0 & 0 \\ g_U\, s_C\, (1-\phi_t) & s_C\, (1-\phi_t) & s_C\, (1-\phi_t) \\ g_U\, s_E\, \phi_t & s_E\, \phi_t & s_E\, \phi_t \end{bmatrix} \end{array},$$

where $g_U$ is the proportion of juvenile biomass maturing to adults in a season, $s_i$ is the proportion of stage $i$ surviving the season, and $\phi_t$ is the proportion of urchin biomass exposed in the open, grazing adult standing kelp in a given season $t$ (the behavior switching function). We explain these parameters further in the following paragraphs.

Growth and maturation is calculated as

$$g_U = \frac{1}{(4 * g_{mat})},$$

reflecting the probability that a juvenile will mature that season given the average age of urchin maturation ($g_{mat}$).

Urchin survival ($s_i$) is a function of natural survival rates ($M_i$) for juveniles and adults, and the rate of predation per kg of predators ($P_i$) for adults, such that

$$s_J = e^{-M_J}$$

$$s_C = e^{-M_H - P_C N_t}, \text{ and}$$

$$s_E = e^{-M_E - \psi P_E N_t}, \psi = \begin{cases} 0, & \sum_{T=1}^{2}(k_{J,t-T} + k_{A,t-T}) < k_{JA}^{min} \\ 1 \end{cases}.$$

Outputs from the Sheephead predator sub-model provided the input values for $N_t$. Here we are assuming linear (Type-I) predation of urchins, where per kg predation by Sheephead is constant. This reasonable for Sheephead, as urchins only make up a small proportion of Sheephead diets and they would be unlikely to reach high, saturating feeding rates (Cowen 1983). However a Type-II or Type-III functional response may be more appropriate for other predators such as rock lobsters (Dunn and Hovel 2020). We also assume that Sheephead do not predate juvenile urchins as juveniles are typically afforded protection by hiding in the spines of adults (Tegner and Dayton 1981). Finally, $\psi$, is a step function that turns off the predation of urchins when a kelp barren state (defined as standing kelp biomass less than a minimum threshold, $k_{JA}^{min}$) has existed for great than 3 months. This captures the active avoidance by Sheephead of starving barren-state urchins (Eurich et al. 2014), which are nutritionally poor and adhere strongly to the substrate (Liebergesell 2022).

The urchin behavioral feeding switch function ($\phi_t$) captures the shift between urchins cryptically grazing on drift kelp when resources are plentiful, to leaving rocky crevices and targeting standing kelp when drift resources are relatively low (Ebeling et al. 1985, Kriegisch et al. 2019, Randell 2022, Rennick et al. 2022a). Using a declining logistic function to model the proportion of urchins grazing upon kelp as a function of drift density, Randell (2022) demonstrated how this behavioral switching can drive alternative kelp-forested and urchin-barren states. We use the same functional form as Randell (2022) but normalize drift density to the total consumptive capacity of urchins (total grazing capacity of the urchin biomass; Rennick et al. 2022a) in a given season. Consequently, the inflection point beyond which more urchins graze standing kelp than drift is when the consumptive capacity exceeds drift supply (i.e., grazing > drift; Figure S1.1). In doing this, we allow for the influence of changing grazing rates with marine heatwaves on urchin switching behavior. A such, the proportion of urchins exposed, grazing standing kelp ($\phi_t$) at time *t* is

$$\phi_t = e^{-v_1\left(\frac{\overbrace{k_{D,t}}^{\text{drift kelp density}}}{\underbrace{p_{D,C}\,(u_{C,t}+u_{E,t})}_{\text{urchin consumptive capacity}}}\right)^{v_2}},$$

where the following are satisfied:

$$v_1 = \frac{v_2 - 1}{v_2} \text{ and } (v_2 - 1)e^{\frac{1-v_2}{v_2}} = w,$$

$w$ is the slope around the inflection point, and $p_{D,C}$ is the maximum grazing rates of drift kelp by cryptic urchins (see below).

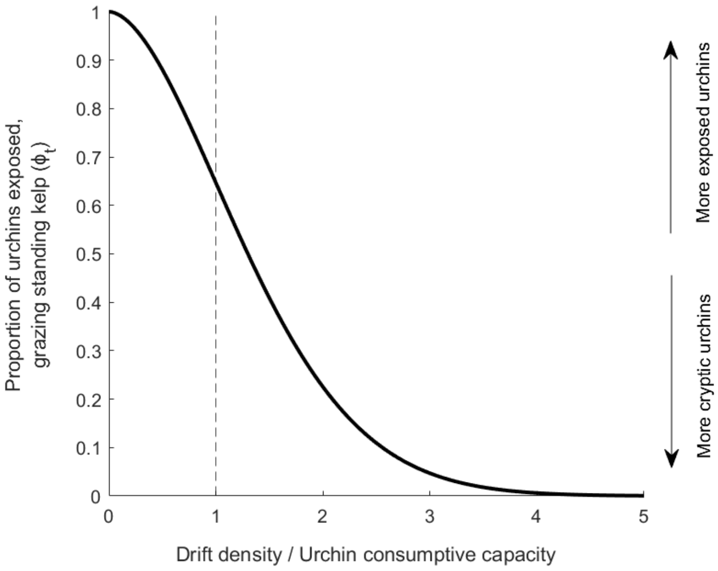


**Figure S1.1** urchin behavioral feeding switch function ($\phi_t$). As the availability of drift kelp to urchin consumptive capacity (biomass of urchins x urchin grazing rate) decreases (left-to-right along the x-axis) more urchins shift to grazing standing kelp. The rate of change of this shift is determined by the slope of the curve ($w_2$), here $w_2 = 0.5$.

*1.4. Urchin-kelp sub-model: giant kelp*

Following the same stage-based matrix approach, the biomass density of kelp in the next time-step ($\mathbf{K}_{t+1}$) is a product of the biomass of urchins in the current time step and the transition matrix ($\mathbf{M}_\mathbf{K}$), such that

$$\mathbf{K}_{t+1} = \mathbf{M}_{\mathbf{K,t}} * \begin{bmatrix} k_{J,t} \\ k_{A,t} \\ k_{D,t} \end{bmatrix},$$

where *J*, *A*, and *D* are the kelp life-stages: standing juvenile sporophytes, standing adult sporophytes, and detached kelp fronds ('drift kelp'), respectively. We define juvenile kelp as sporophytes that are visible to the naked eye, but not mature (0-3 months old; Schiel and Foster 2015). Juveniles are considered standing kelp and are, therefore, grazed by urchins and contribute to the production of drift.

The stage transition matrix ($\mathbf{M}_\mathbf{K}$) in timestep *t* for kelp is

$$\mathbf{M}_{\mathbf{K,t}} = \begin{matrix} \\ juvenile \\ adult \\ drift \end{matrix} \begin{matrix} juvenile & adult & drift \\ \left[\begin{matrix} 0 \\ g_K\,\gamma\,r_S\,\kappa_{A,E}\,(1-c) \\ c\,r_D\,\kappa_{D,H} \end{matrix}\right. & \begin{matrix} R_K\,s_{Y,t}\,\theta_{K,t}\,\gamma\,r_S\,\kappa_{Y,E} \\ g_K\,\gamma\,r_S\,\kappa_{A,E}\,(1-c) \\ c\,r_D\,\kappa_{D,H} \end{matrix} & \left.\begin{matrix} 0 \\ 0 \\ (1-d)\,r_D\,\kappa_{D,H} \end{matrix}\right] \end{matrix}.$$

We explain all parameters in the following paragraphs (see also Table S1.3).

As giant kelp spores have short dispersal distances (generally <1 km; Gaylord et al. 2002, Reed et al. 2004, Schiel and Foster 2015), we assume closed-patch dynamics, where the total incoming spore biomass is a function of local adult standing kelp biomass densities. The biomass density of settled spores per biomass of adults ($R_K$) includes zoospore production, survival and development through the gametophyte phase, successful fertilization, and settlement survival. To account for interannual variation in settling spores:

$$R_K = norm(\overline{R_K}, \sigma_{RK}),$$

where $\overline{R_K}$ is yearly mean biomass and $\sigma_{RK}$ is the yearly variance of incoming kelp spores.

In southern California, peaks in recruited juvenile sporophytes occur summer to autumn (July-November; Reed et al. 2008a), following winter peaks in spore production (Reed et al. 1996). Capturing this seasonal variation, $\theta_{K,t}$ (the kelp recruitment timing function) distributes recruitment proportionally over the seasons and sums to one each year. Therefore, the total number of incoming kelp recruits in timestep *t* is

$$Y_t = R_k\,\theta_{K,t}\,k_{J,t},$$

Incoming kelp recruits compete for space and are negatively affected by shading of older standing adult kelp biomass (Nisbet and Bence 1989, Reed 1990, Stewart et al. 2009). As such, we used a combined Ricker and Beverton-Holt function, which captures both inter- and intra-cohort density dependent effects (White 2009). Per biomass recruit survival ($s_Y$) is, therefore,

$$\bar{s}_{Y,t} = \frac{(D-1)\, k_{A,t}\, e^{\beta\, D\, k_{A,t}}}{D\, Y_t\, e^{\beta\, D\, k_{A,t}} + e^{\beta\, k_{A,t}}\, ((D-1)\, k_{A,t} - D\, Y_t)},$$

where $\beta$ is the strength of density dependence and D is the relative strength of inter- versus intra-cohort density-dependent processes (0 < D < 1). Since D is unknown, we considered the two extremes relevant for giant kelp: first, D is relatively low (D = 0.01) reflecting the case where the effect of a kilogram of adults on recruit survival is far greater than that of a kilogram of recruits. In this case, the effects of shading (inter-cohort density dependence) are dominate. As adults are unlikely to have less than an equal effect on recruit survival than other recruiting sporophytes, the second extreme we consider is where adults and recruits have equal effect (D = 0.5). Our main results were not overly sensitive to the value of D, and we present only the results where the effects of shading are dominate (D = 0.01; see also Section 2: Sensitivity Analysis).

Since kelp dynamics occur at a smaller scale than our 1 ha model domain (i.e., at the local transect scale), we approximated the model landscape scale density dependent survival (following White 2011) as

$$s_{Y,t} \approx \bar{s}_{Y,t} + 0.5\, \bar{s}''_{Y,t} + var(k_{A,t})$$

where $var(k_{A,t})$ is the spatial variance in the density of adult standing kelp across the model unit space.

Reflecting fast maturation rates (on the order of months; Schiel and Foster 2015), we assumed that juvenile sporophytes become adults, or contribute to drift, in the next season. Adult sporophyte growth over time is captured through $g_K$, the seasonal increase in standing kelp biomass, and $\gamma$, the relative change in standing (juvenile and adult) kelp biomass over the seasons.

Non-grazing loss of standing kelp biomass is captured in two terms: *c*, proportion of juvenile and adult standing kelp biomass converted to drift kelp in a time step, and $r_S$, the proportion of non-grazed standing kelp biomass that survives the season. $r_S$ accounts for storm and wave damage, senescing blade material, boat propeller damage, and dissolved material released from blades and stipes (Rassweiler et al. 2021). Drift kelp biomass is either lost through decomposition (*d*) or removed from the system via processes such as transportation through wave action or storm events, where $r_D$ is proportion of non-decomposed drift biomass retained in system.

We assumed Type-II grazing functional response for standing and drift kelp by urchins, following Randell (2022). This is reasonable given that, like most herbivores following a Type-II saturating function, urchins require handling time between kelps stipes (Boada et al. 2017). $\kappa_{i,j}$ is the proportion of kelp stage *i* (juvenile, adult, or drift), grazed by urchins stage *j* (juvenile, cryptic adult, exposed adult), such that:

$$\kappa_{i,j} = e^{-\Omega_{i,j}} ,$$

where the instantaneous rate of grazing $-\Omega_{i,j}$ of kelp stage *i*, by urchins stage *j*, is a per-kilogram Type-II response :

$$\Omega_{i,j} = \frac{a_{i,j,t}}{1+\frac{1}{p_{i,j}} a_{i,j,t} k_{i,t}} u_{j,t},\ \ a_{i,t} > 0 \ and \ p_i > 0.$$

$a_{i,j}$ is the per kilogram kelp mortality due to grazing by a kilogram of urchins in the absence of conspecifics (search/attack rate), and $p_{i,j,t}$ is the asymptotic (max) biomass of prey consumed by an urchin in a time step (1/handling and ingestion time), in season *t*. This assumes that there is no effect of increasing urchin densities (Rennick et al. 2022a). Note that exposed urchins eat recruits and adult standing kelp, and cryptic urchins only eat drift kelp:

$$\boldsymbol{a} = \begin{array}{r} \\ \text{juvs} \\ \text{adult} \\ \text{drift} \end{array} \begin{array}{c} \begin{array}{cc} \text{cryptic} & \text{exposed} \end{array} \\ \begin{bmatrix} 0 & a_{J,E} \\ 0 & a_{A,E} \\ a_{D,C} & 0 \end{bmatrix} \end{array} \qquad \boldsymbol{p}_{,t} = \begin{array}{r} \\ \text{juvs} \\ \text{adult} \\ \text{drift} \end{array} \begin{array}{c} \begin{array}{cc} \text{cryptic} & \text{exposed} \end{array} \\ \begin{bmatrix} 0 & p_{J,E,t} \\ 0 & p_{A,E,t} \\ p_{D,C,t} & 0 \end{bmatrix} \end{array}$$

**Table S1.2** State variables for the urchin-kelp sub-model. Each state variable is the biomass density (per ha) at season *t*.

| | | |
|---|---|---|
| *Urchins* | $u_{J,t}$ | Juvenile (small, sexually immature) urchins |
| | $u_{C,t}$ | Cryptic adult urchins |
| | $u_{E,t}$ | Exposed adult urchins |
| *Kelp* | $k_{J,t}$ | Juvenile standing kelp |
| | $k_{A,t}$ | Adult standing kelp |
| | $k_{D,t}$ | Drift kelp |

**Table S1.3** Parameter symbols, descriptions, values and references (where applicable) for the urchin-kelp sub-model. Values in red indicate values used during heatwave periods, and their associated references, where applicable. Unless otherwise specified, urchin data is for purple urchins (*Strongylocentrotus purpuratus*) and kelp data is for giant kelp (*Macrocystis pyrifera*; hereafter 'kelp'). Note that all relevant values are per hectare ($ha^{-1}$). Conversions used for urchin abundance to wet biomass were: 1) for size to biomass, Biomass (kg) = 0.0499 x test dimeter (cm) + 0.0019 (Ling et al. 2015), and 2) for number to biomass, Average purple urchin weight = 0.21kg (PISCO data[*]). Conversions used for kelp stipe numbers to wet biomass were: 1) for number to dry biomass, dry weight = 0.33stipes + 0.16 (Reed et al. 2009), and 2) from dry to wet biomass, wet-to-dry ratio = 0.094 (Rassweiler et al. 2018).

| | Symbol | Description | Baseline value *(Heatwave value)* | Units | References | Notes |
|---|---|---|---|---|---|---|
| **Urchins** | $\overline{R_U}$ | Mean larval production (fecundity), dispersal and settlement for urchins (assumes open population) | $5x10^5$ | $kg.yr^{-1}$ | | Value set to achieve a range of urchin biomasses that fall within observed values for Channel Islands kelp forests, based on PISCO data[*]. Figures S1.2-S1.3 |
| | $\sigma_{RU}$ | Variance of incoming recruits | 0.621 | | | Normalised variance from PISCO data[*] for central and northern California. No urchin recruit data exists for Channel Islands. |
| | $\theta_{U,t}$ | Urchin recruitment timing function | [0.05 0.54 0.36 0.05][+] | | Okamoto et al. (2020b) | |
| | $g_{mat}$ | Average urchin maturation age | 2 | yrs | Gonor (1972) | |
| | $s_i$ | Proportion of urchin biomass stage *i* surviving the season | | | | See main text for relevant functions |
| | $M_j$ | Instantaneous natural mortality rate of stage *j* (not including predation) | $\begin{matrix} juvs \\ cryptic \\ exposed \end{matrix} \begin{bmatrix} 0.1 \\ 0.1 \\ 0.1 \end{bmatrix}$ | $season^{-1}$ | Russell (1987), Dunn et al. (2017) | |
| | $P_i$ | Instantaneous predation mortality rate of stage *j* (per kg predator) | $\begin{matrix} cryptic \\ exposed \end{matrix} \begin{bmatrix} 0.0065 \\ 0.013 \end{bmatrix}$ | $kg\ pred^{-1} .season^{-1}$ | Nichols et al. (2015), Selden et al. (2017b) | |
| | PLD | Urchin planktonic larval duration | 65 | days | Arafeh-Dalmau et al. (2022) | |
| | $w$ | Slope around inflection point of behaviour switching function | 0.5 | | | Value set so that kelp forest switches to urchin barren around observed urchin biomass thresholds (Ling et al. 2015). Figure S1.2 |
| | $\psi$ | Step function turning off predation of urchins in a kelp barren state | | | | See main text for relevant functions |
| | $k_{J+A}^{min}$ | Standing kelp density threshold below which a kelp barren state is declared | 1170 | kg | | Set to 1% of the average standing kelp densities observed at a persistent kelp forest location (Anacapa East) in the Channel Islands (~700kg.60m2/0.006 = ~ $1.17*10^5$kg.ha), based on PISCO data[*]. |

| | Symbol | Description | Baseline value *(Heatwave value)* | Units | References | Notes |
|---|---|---|---|---|---|---|
| **Kelp** | $\overline{R_K}$ | Yearly mean zoospore production, successful fertilization and settlement of spore, *per kg standing kelp* | $4x10^4$ ***(baseline x ½)*** | kg.year$^{-1}$ | Dayton et al. (1984), Schiel and Foster (2015), Hollarsmith et al. (2020) | Value calculated as follows:<br>• ~$10^{11}$ zoospores per plant (Schiel and Foster 2015, pg. 31)<br>• Halve value due to 50:50 sex ratio (Graham 2007, pg. 47)<br>• ~1% survival probability to settled microsporophytes (Schiel and Foster 2015, pg. 77)<br>• ~20% survival probability for settled spores lasting 3 months (Dayton et al. 1984)<br>∴ $10^{11}$ x 0.5 x 0.01 x 0.2 = ~$10^8$ spores per plant<br>• We assumed that one recruit weighs, on average, 1g<br>• One adult plant = ~10kg wet (Reed et al. 2009, Rassweiler et al. 2018)<br>∴ ~$10^8$ spores per stipe = $10^8$ x 0.001 x 0.1<br>= $10^4$ kg recruits per kg adult (*per season*)<br><br>*Note that Schiel & Foster (2015, Fig 4.2A inset) have initial settlement rates on the order of $10^6$-$10^8$ spores.m$^2$, which converts to $10^2$-$10^4$ kg of spores per kg adult stipes.* |
| | $\sigma_{RK}$ | Variance of incoming settlers | 0.389 | | | Normalised variance from PISCO data* for Channel Islands California. |
| | $\beta$ | Strength of density dependence effect of shading by adults | $9x10^{-5}$ | | | Value set to achieve a range of standing kelp biomasses that fall within observed values for Channel Islands kelp forests, based on PISCO data*. Figures S1.2-S1.3.<br>Sensitivity analysis value: $1x10^{-5}$ (paired with D = 0.5) |
| | $D$ | Relative strength of inter- versus intra-cohort density-dependent processes | 0.01 | | White (2009) | Sensitivity analysis value: 0.5 (paired with $\beta$ = $1x10^{-5}$) |
| | $var(\overline{k_{J+A,t}})$ | Variance in the density of standing kelp across the model unit space | 189,090 | | | Spatial variance at transect level, averaged over years, based on PISCO* kelp surveys at the Channel Islands. |
| | $\theta_{K,t}$ | Kelp recruitment timing function | [0.1, 0.1, 0.4, 0.4]$^+$ | | Reed et al. (2008b) | |
| | $r_S$ | Proportion of non-grazed standing kelp biomass remaining at the end of the season | 0.5688 | season$^{-1}$ | Rassweiler et al. (2018) | Conversion from average daily plant loss rate to proportion surviving a season (91 days): exp(-0.0062*91) = 0.5688.<br>We assumed that biomass is proportional to number of plants. |
| | $g_K$ | Average increase in adult kelp biomass as it ages | 6.825 | kg.season$^{-1}$ | Stewart et al. (2009) | Conversion from average daily growth rate to seasonal (91 days): 0.075 x 91 = 6.825kg fronds per season |

| | Symbol | Description | Baseline value<br>***(Heatwave value)*** | Units | References | Notes |
|---|---|---|---|---|---|---|
| | $\gamma$ | Relative change in standing kelp biomass over the seasons | [1, 1, 0.8, 0.9] [+]<br>***([1, 1, 0.5, 0.5])*** | | Zimmerman and Kremer (1986) | Based on average, pre-heatwave water temperatures observed in the Channel Islands. |
| | $c$ | Proportion of standing kelp biomass converting to drift (*drift production*) | 0.9 | season$^{-1}$ | Rassweiler et al. (2018), Rennick et al. (2022b) | Conversion from daily to seasonal proportion of biomass converted to drift = 1-(1-0.025)$^{91}$. |
| | $r_D$ | Proportion of drift biomass retained locally | 0.7 | season$^{-1}$ | Hobday (2000), Figurski (2010) | |
| | $d$ | Proportion of drift decomposing | 0.1 | season$^{-1}$ | Figurski (2010) | |
| | $a_{i,j}$ | Per kg mortality of kelp stage $i$ due to grazing by an urchin, stage $j$, in the absence of conspecifics (*search/attack rate*) | $\begin{array}{l} \\ juv\ k \\ adult \\ drift \end{array} \begin{array}{c} hiding \quad exposed \\ \begin{bmatrix} 0 & 0.5 \\ 0 & 0.5 \\ 1 & 0 \end{bmatrix} \end{array}$ | kg kelp .kg urchin$^{-1}$ .season$^{-1}$ | Kriegisch et al. (2019) | No data is available to parameterise this, so we assumed very high attack rates with near-zero standing or drift densities. Kriegisch et al. (2019) findings suggest that attack rates for standing are less than that for drift as kelp has lower consumption than drift at high resource density (kelp forests). |
| | $p_{i,j,t}$ | Max kg of kelp stage $i$ consumed by an urchin, stage $j$, in a time step (1/*handling time*) | $\begin{array}{l} \\ juv\ k \\ adult \\ drift \end{array} \begin{array}{c} hiding \quad exposed \\ \begin{bmatrix} 0 & p_t \\ 0 & p_t \\ p_t & 0 \end{bmatrix} \end{array}$<br><br>$p_t = 2.985 * [1\ 0.9\ 1.15\ 1.2]$ [+]<br><br>$(p_t = 2.985 * [1.15\ 1.05\ 1.2\ 1.3])$ | kg kelp .kg urchin$^{-1}$ .season$^{-1}$ | Foster et al. (2015), Rennick et al. (2022b) Spindel (2023) | Kriegisch et al. (2019) findings suggest that the maximum feeding rate is the same for drift and standing kelp, as consumption rates were the same for drift and kelp in urchin barrens (when resources are low). We assume they are the same. Experimental data (Randell 2022, Rennick et al. 2022b) have a range of maximum feeding rate values and we used the average of these to obtain the baseline value.<br>• Rennick et al. (2022): 0.02 g detritus.g urchins$^{-1}$.day$^{-1}$.<br>Conversion = 0.02*1000/1000*91<br>= 1.82 kg kelp.kg urchin$^{-1}$.season$^{-1}$.<br>Urchins were taken from a kelp forest and fed for a week prior to experiments. Experiments were run at 14.5°C water temperature.<br>• Foster et al (2015): 1.14 g kelp.urchin$^{-1}$.day$^{-1}$.<br>Conversion = 1.14/1000*91<br>= 0.104 kg kelp.urchin$^{-1}$.season$^{-1}$<br>= 0.104/0.025<br>(mean experimental urchin weighed 25g)<br>= 4.15 kg kelp.kg urchin$^{-1}$.season$^{-1}$<br>Urchins were taken from an established urchin barren and starved for a week prior to experiments. Experiments were run at 11.6 to 16.3°C water temperature.<br>Finally, we used temperature experiments from Spindel (2023) to estimate relative change in max feeding rates over the seasons (based on average, pre-heatwave water temperatures observed in the Channel Islands.). |

[*] See PISCO_kelp-urchin-Sheephead_data.Rmd file for details. [+] Seasonal variations, in the order winter, spring, summer, autumn

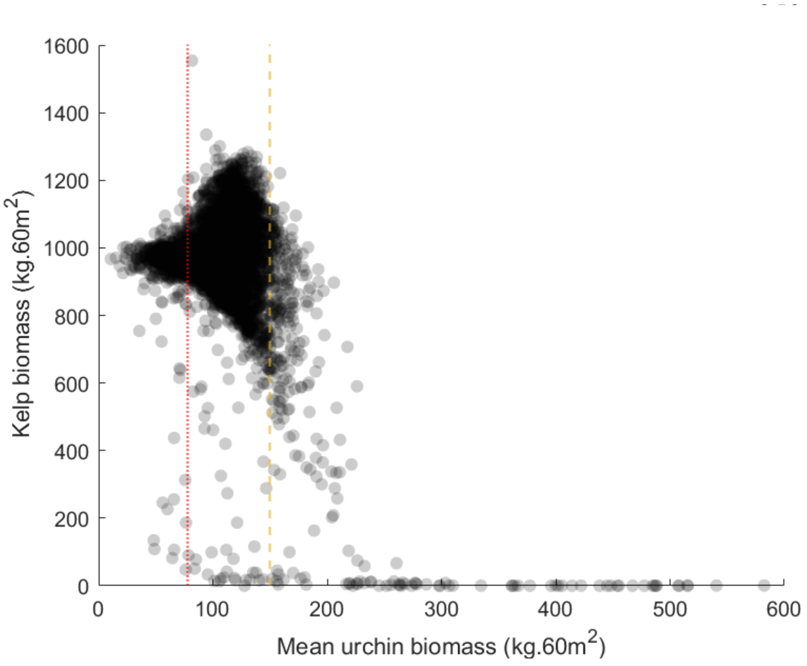


**Figure S1.2** Example average standing kelp and urchin biomass densities in the final year of the model run. Each point is an individual model simulation (n=10,000). Vertical lines indicates approximate Californian (red dotted line) and global (orange dashed line) threshold urchin biomass densities, above which kelp forests have shifted to urchin barrens (Ling et al. 2015).

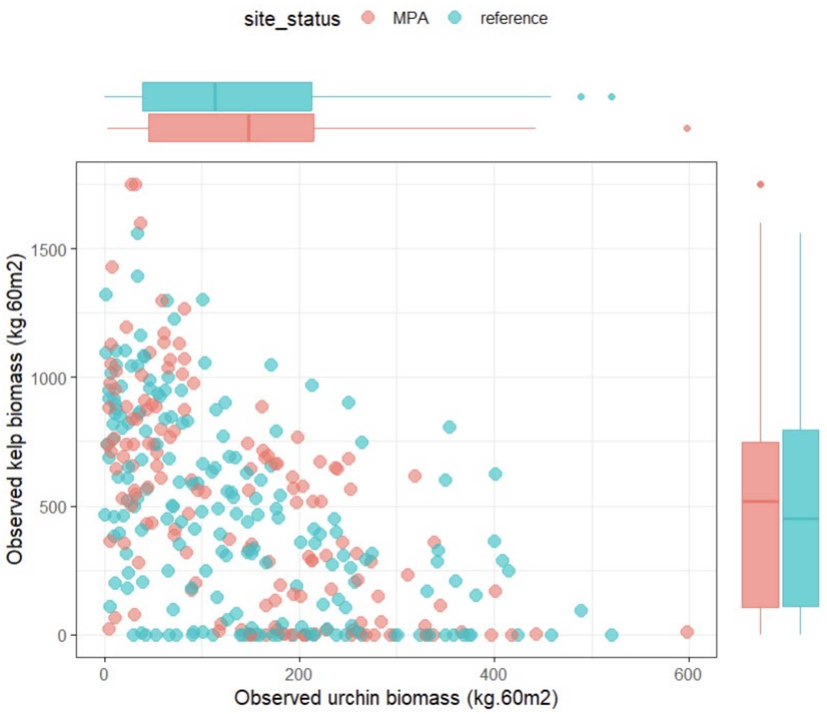


**Figure S1.3** Site-averaged standing kelp and urchin biomass densities in the Channel Islands, California.

#### 2. Sensitivity analyses

We conducted sensitivity analyses to determine the relative importance of changes in model parameters to our main results. We first undertook a **global sensitivity analysis** following the approach of Harper et al. (2011), which uses regression trees to rank and assign relative importance to parameters. In short, we ran $10^4$ solutions of the model, each randomly drawing a selection of parameter values from biologically reasonable ranges. Note that each solution had $10^5$ replicates, as per the baseline runs. Using the *proportion of model simulations with persisting kelp forests over time* as our response variable, the relative importance value of each parameter was determined using a random forest procedure, with higher values indicating greater sensitivity to that model parameter (Table S2.1). We ran the model under the baseline scenario, without the heatwave or interventions.

We then undertook a **local sensitivity analysis** to evaluate the importance of changes in the top 5 globally important parameters, as identified in the global sensitivity analysis (Table S2.1). In the local sensitivity analysis, we ran the model under low, baseline, and high values for each parameter (Table S2.1), while setting all other parameters to their baseline values. We focused on the case where the three individual management scenarios (fishery closure, urchin removal, and kelp seeding) lasted three years, starting one year before the heatwave, concurrent with the heatwave, and one year after the start of the heatwave. Also, we focused on the most intensive management scenarios in which 100% of the pre-heatwave urchin and kelp biomass is removed or seeded each year, respectively. Here, we used *the percentage increase in probability of kelp forest persistence* as our response variable.

We found that our overall results – that fishery closures were most effective before, while urchin removal and kelp seeding were most effective during a heatwave, and that kelp seeding is notably less effective that other interventions – were mostly robust to changes in key parameter values (Figure S2.1). Of note was when predator densities ($N_t$) or predation rates ($P_C$) were high, and when predator densities ($N_t$) were low (stars in Figure S2.1). Decreasing predation densities ($N_t$) increased the relative effectiveness of fishery closures and urchin removal, and decreased the effectiveness of kelp seeding, and vice versa for increasing predation densities. This makes intuitive sense as changing local predator density shifts the balance between top-down and bottom-up processes. Increasing predator densities strengths top-down controls, decreasing the effectiveness of fishery closures or urchin removal, but increasing the effectiveness of kelp seeding. Likewise, increasing predation rates ($P_C$), exerts greater top-down controls, again increasing the effectiveness of kelp seeding, but also resulting in fishery closures being consistently more effective than urchin culling throughout the heatwave.

**Table S2.1** Parameter importance as ranked by the *global sensitivity analysis*, and the low, baseline, and high parameter values explored in the local sensitivity analysis (LSA). Bold parameters indicates the top-5 parameters explored in the LSA.

| Parameter | Name | Importance | LSA Range *[low, baseline, high]* |
|---|---|---|---|
| $P_C$ | Cryptic urchin predation mortality | 213.91 | [0.0032, 0.0065, 0.013] |
| $r_S$ | Standing kelp survival (not including grazing) | 125.66 | [0.75, 0.9, 0.95] |
| $c$ | Standing kelp to drift conversion rate | 119.18 | [0.25, 0.5688, 1] |
| $g_K$ | Kelp growth | 93.45 | [3.4125, 6.825, 13.65] |
| $N_t$ | Biomass of urchin-predating Sheephead | 47.18 | [50*, 180*, 360*] |
| $w$ | Behaviour switching function shape parameter | 2.25 | |
| $var(\overline{k_{J+A,t}})$ | Standing kelp spatial variance | 2.22 | |
| $M_C$ | Cryptic urchin natural mortality (not including predation) | 1.77 | |
| $p_t$ | Max kg of kelp grazed by urchins | 1.28 | |
| $\overline{R_U}$ | Mean biomass of urchin recruits | 1.22 | |
| $\sigma_{RK}$ | Incoming kelp recruit variance | 1.20 | |
| $a_{D,C}$ | Search/attack rate on drift kelp by cryptic urchins | 0.79 | |
| $M_E$ | Exposed urchin natural mortality (not including predation) | 0.39 | |
| $d$ | Drift kelp decomposition | 0.27 | |
| $r_D$ | Drift kelp retention | 0.20 | |
| $M_J$ | Juvenile urchin natural mortality (not including predation) | 0.12 | |
| $\beta$ | Strength of density-dependent kelp recruitment | -0.01 | |
| $g_{mat}$ | Average urchin maturation age | -0.31 | |
| D | relative strength of inter- versus intra-cohort density-dependent processes | -0.58 | |
| $P_E$ | Exposed urchin predation mortality | -0.70 | |
| $a_{J,E}$ , $a_{A,E}$ | Grazing mortality of juvenile and adult kelp by exposed urchins | -0.91 | |
| $k_{J+A}^{min}$ | Barren state threshold | -0.98 | |
| PLD | Urchin planktonic larval duration | -1.38 | |
| $\overline{R_K}$ | Mean biomass of kelp recruits | -2.19 | |
| $\sigma_{RU}$ | Incoming urchin recruit variance | -2.96 | |

* Time-averaged, mean Sheephead biomass across all replicates, for the baseline (fished) scenario. This value is set by varying the density of incoming Sheephead recruits ($\overline{R_U}$).

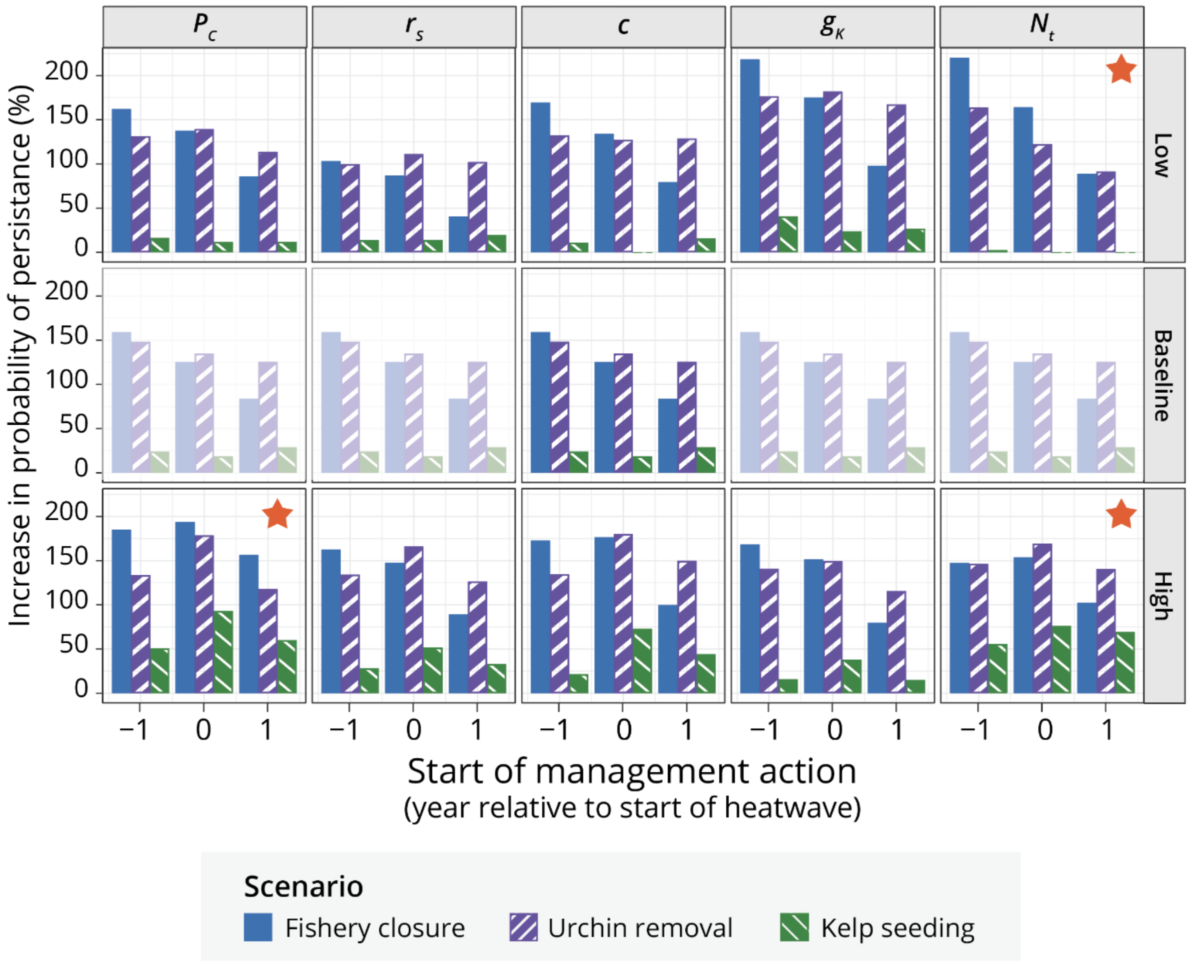


**Figure S2.1 Local sensitivity analysis**. Percentage increase in the probability of kelp forest persistence through a 2-year marine heatwave with individual management actions, over timing of action and changes in parameter values (rows). Columns are parameters (see Tables S3.1 & S2.1). Note that the middle row is the same plot showing baseline values. Starred facets indicate notable deviations from baseline results.

In addition to evaluating the sensitivity of the model outcomes to parameter values, we also considered the sensitivity of results to 1) the relative strength of inter- versus intra-cohort density-dependent processes (D) and 2) the sampling time.

First, our main results were not overly sensitive to the value of D, but intervention actions overall were less effective when inter- and intra-cohort density dependence was of equal strength (D = 0.5; Figure S2.2).

Second, in the main text we sampled model runs 8 years after the heatwave had finished, enough time for all mitigation actions to finish. Here, we considered an alternative sampling time that was either 1 year after the heatwave or when the mitigation action ended, whichever was last. We found that monitoring sooner only reduced the realized effects of temporary fishery closures (Figure S2.3); that is the effects of temporary fishery closures are lagged due to the time required to build-up predator biomass (Figure 2, main text).

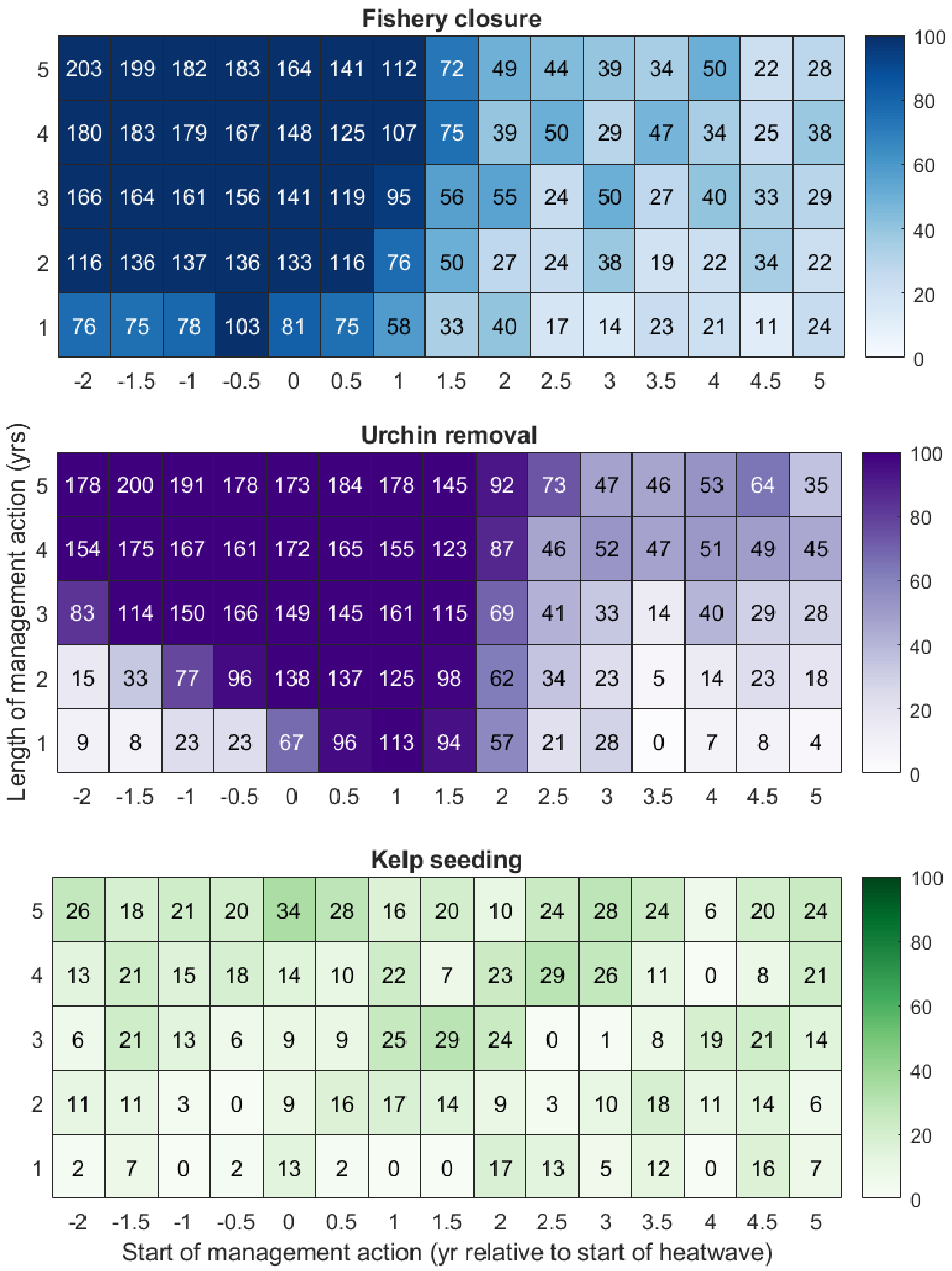


**Figure S2.2 Sensitivity to the relative strength of inter- versus intra-cohort density dependence (D).** Percentage increase in the probability of kelp forest persistence through a 2-year marine heatwave with individual management actions, over length and timing of action. These modelled scenarios present the most intensive management scenario in which 100% of the pre-heatwave urchin and kelp biomass is removed or seeded each year, respectively. See also Figure 3 in main text.

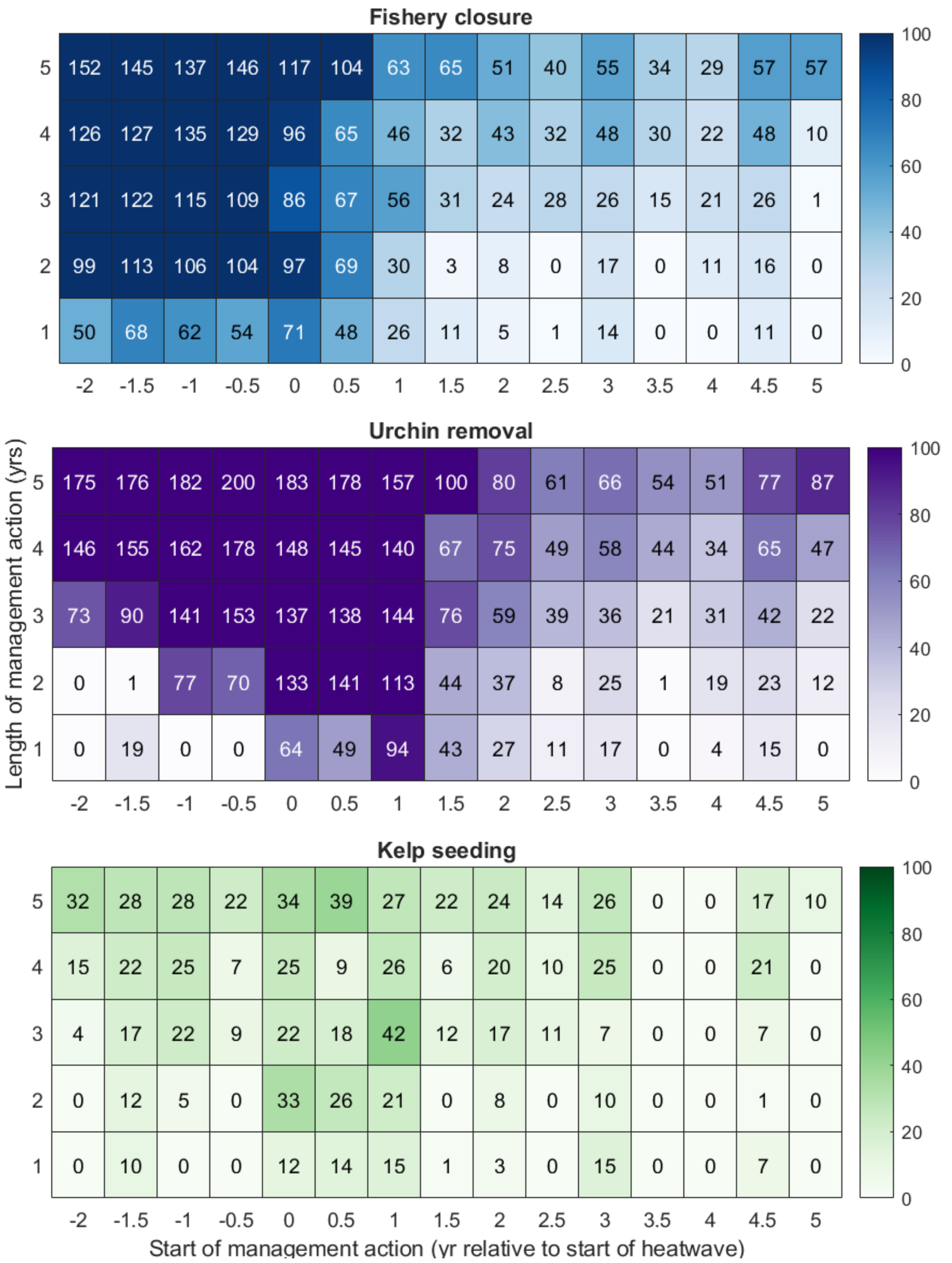


**Figure S2.3 Sensitivity to sampling time**. Results when sampled either 1 year after the heatwave or the mitigation action ended, whichever was last: Percentage increase in the probability of kelp forest persistence through a 2-year marine heatwave with individual management actions, over length and timing of action. These modelled scenarios present the most intensive management scenario in which 100% of the pre-heatwave urchin and kelp biomass is removed or seeded each year, respectively. See also Figure 3 in main text.
